# A dual, catalytic role for the fission yeast Ccr4-Not complex in gene silencing and heterochromatin spreading

**DOI:** 10.1101/2023.03.27.534317

**Authors:** Drice Challal, Alexandra Menant, Can Goksal, Estelle Leroy, Bassem Al-Sady, Mathieu Rougemaille

## Abstract

Heterochromatic gene silencing relies on combinatorial control by specific histone modifications, the occurrence of transcription, and/or RNA degradation. Once nucleated, heterochromatin propagates within defined chromosomal regions and is maintained throughout cell divisions to warrant proper genome expression and integrity. The fission yeast Ccr4-Not complex has been involved in gene silencing, but its relative contribution to distinct heterochromatin domains and its role in nucleation versus spreading have remained elusive. Here, we unveil major functions for Ccr4-Not in silencing and heterochromatin spreading at the mating type locus and subtelomeres. Mutations of the catalytic subunits Caf1 or Mot2, involved in RNA deadenylation and protein ubiquitinylation respectively, result in impaired propagation of H3K9me3 and massive accumulation of nucleation-distal heterochromatic transcripts. Both silencing and spreading defects are suppressed upon disruption of the heterochromatin antagonizing factor Epe1. Overall, our results position the Ccr4-Not complex as a critical, dual regulator of heterochromatic gene silencing and spreading.

**Author Summary:** Eukaryotic genomes are partitioned into relaxed, gene-rich regions, and condensed, gene-poor domains called heterochromatin. The maintenance of heterochromatin is crucial for proper genome expression and integrity, and requires multiple factors regulating histone modifications and/or the levels of RNA molecules produced from these regions. Such effectors not only promote heterochromatin assembly but also ensure its propagation from specific nucleation sites to defined domain boundaries. However, while the mechanisms involved in initiation of heterochromatin formation have been well documented, the molecular and biochemical properties underlying its spreading remain largely elusive. By combining genetic and single-cell approaches, we report here that the fission yeast Ccr4-Not complex, a multisubunit complex conserved throughout eukaryotes, is essential for efficient heterochromatin spreading to repress expression of nucleation-distal RNAs. The two catalytic activities of the complex, RNA deadenylation and protein ubiquitinylation, are each critical, thereby defining a dual enzymatic requirement in the process.

## Introduction

Eukaryotic genomes organize into gene-rich, euchromatic regions and transcriptionally-repressed heterochromatin domains. The assembly, maintenance and inheritance of heterochromatin is essential for major biological processes, including gene expression, chromosome segregation, genome stability and cell fate [1,2]. In the fission yeast *Schizosaccharomyces pombe* (*S. pombe*), heterochromatin assembles at defined chromosomal regions, including pericentromeric repeats, subtelomeric regions and the silent mating type locus. These domains are enriched in nucleosomes methylated on histone H3 lysine 9 (H3K9me), a modification catalyzed by the sole Suv39 homolog Clr4 and bound by proteins of the HP1 family [1]. Such structural components constitute a platform for the recruitment of additional silencing and heterochromatin assembly complexes, including histone-remodeling and modifying enzymes as well as the RNA interference (RNAi) machinery [3-6]. These different effectors tightly cooperate in intricate, often inter-dependent regulatory pathways to mediate transcriptional (TGS) and co- or *cis* post-transcriptional (C/*cis*-PTGS) gene silencing, which ultimately restrict RNA polymerase II (RNAPII) accessibility and the accumulation of heterochromatic transcripts, respectively. Heterochromatin also engages factors endowed with anti-silencing functions, such as the H3K9me antagonizing protein Epe1 [7-12], and the dynamic balance between opposite activities is believed to prevent spreading within adjacent euchromatin and maintain heterochromatin throughout cell divisions [13-17].

Despite commonalities in core components of the silencing machinery, the mechanisms that drive heterochromatin-based silencing substantially differ between loci: while RNAi is essential at pericentromeres, it has little impact at telomeres and the mating type locus due to functional redundancy with RNAi-independent mechanisms. The shelterin complex and the CREB-family proteins Atf1/Pcr1 bind to specific *cis*-acting DNA sequences and independently recruit Clr4 to nucleate heterochromatin at these regions [16,18-24]. The intrinsic capacities of Clr4 to both recognize H3K9me and methylate adjacent nucleosomes define the molecular basis of a “read-write” mechanism by which heterochromatin subsequently propagates throughout entire domains [25-28]. Heterochromatin spreading from nucleation sites requires the Clr4-mediated transition from H3K9me2 to H3K9me3, which is important for the switch to TGS [25,26,29]. Despite these progresses, our understanding of the mechanisms underlying heterochromatin propagation is still far from being complete.

Beyond RNAi, other RNA processing/degradation machineries contribute to heterochromatic gene silencing [30-34]. Among these is the conserved, multifunctional Ccr4-Not complex, which contains two catalytic modules: one formed by the RNA deadenylases Ccr4 and Caf1 and another comprising the E3 ubiquitin ligase subunit Mot2/Not4 [35-38]. Previous studies in *S. pombe* showed that Ccr4-Not mediates deposition of H3K9me2 at rDNA repeats, subtelomeric regions and a subset of meiotic genes [39,40]. The complex was also found to act redundantly with the RNAi machinery to target chromatin-bound RNAs for degradation and preserve heterochromatin integrity [34]. Recently, Ccr4-Not was involved in the control of transcriptional efficiency by limiting the levels of RNAPII-associated heterochromatic transcripts [41]. Though, the precise function and contribution of Ccr4-Not to gene silencing, heterochromatin assembly and spreading remain unclear.

Here, we demonstrate that Ccr4-Not on its own is crucial for heterochromatic gene silencing and spreading. Mechanistically, the deadenylation and ubiquitinylation activities of the complex independently mediate gene repression, maintain heterochromatin integrity and ensure its efficient propagation within the mating type locus and at subtelomeres. We further show that these functions of the complex are antagonized by the jumonji protein Epe1. Together, our findings unveil a dual, catalytic role for Ccr4-Not in gene silencing and heterochromatin spreading, thereby highlighting its biological relevance to preserve genome expression and integrity.

## Results

### The Ccr4-Not complex mediates heterochromatic gene silencing at the mating type locus

To assess the role of the Ccr4-Not complex in heterochromatic gene silencing, we generated strains carrying the *ura4*+ reporter gene inserted at pericentromeric repeats (otr1R::*ura4*+), subtelomeric regions (tel2L::*ura4*+) or the silent mating type cassette (mat3M::*ura4*+) in which non-essential subunits (all but the scaffolding subunit Not1) were deleted (**Fig 1A, 1B**). Gene silencing was probed by growing cells in the presence of 5FOA, which counter-selects those expressing *ura4*+, and by measuring steady state *ura4*+ mRNA levels in RT-qPCR assays. A strain lacking the H3K9 methyltransferase Clr4, defective for silencing at all heterochromatic domains, was assayed in parallel as an internal control. As shown in **Fig 1C, 1D**, we observed a minor defect in pericentromeric silencing in the *caf1*Δ mutant, while *mot2*Δ cells did not accumulate *ura4*+ transcripts despite a partial sensitivity to 5FOA. These two mutants also exhibited a significant increase in *ura4*+ mRNA levels when expressed from subtelomeres, although this was considerably lower than that of cells lacking Clr4 (about 15 to 20-fold compared to 500-fold) (**Fig 1E, 1F**). Consistent with these results and previous observations [34,41], endogenous dg pericentromeric repeats and subtelomeric *tlh1+* sequences were not or only marginally increased in *caf1*Δ and *mot2*Δ mutants (**S1A Fig**), arguing against a major contribution of the Ccr4-Not complex *per se* at these loci.

**Fig 1.**
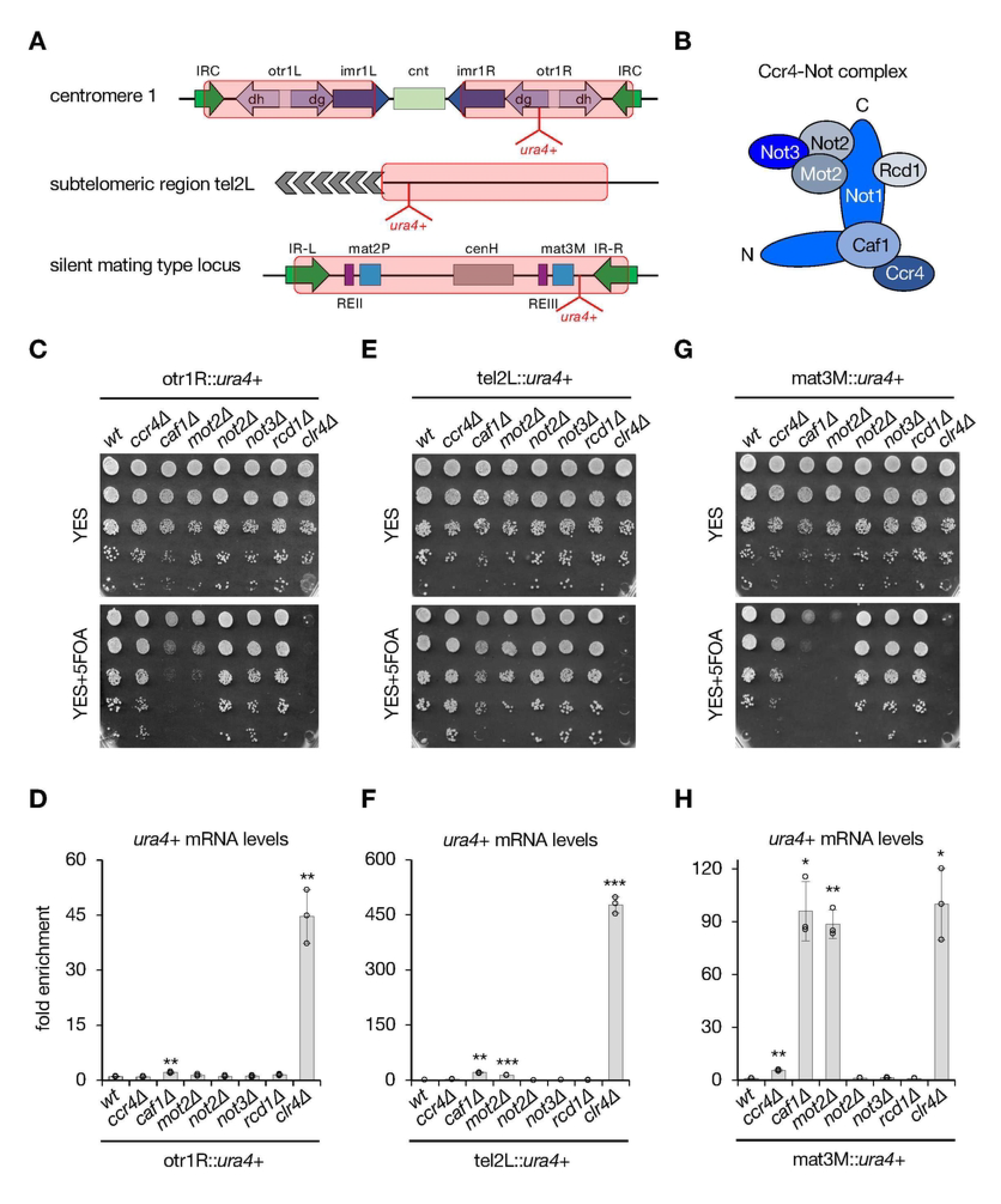
The Ccr4-Not complex mediates heterochromatic gene silencing at the mating type locus. **A.** Scheme of the different heterochromatin domains with the position of the *ura4*+ reporter gene. Red rectangles define heterochromatin regions. **B.** Scheme illustrating the different subunits of the core Ccr4-Not complex. **C.**, **E.**, **G.** 5-fold serial diluted silencing assays using the *ura4*+ reporter gene inserted at pericentromeric repeats (otr1R::*ura4*+), subtelomeric regions (tel2L::*ura4*+) and the mating type locus (mat3M::*ura4*+). Cells of the indicated genotypes were plated on both non-selective (YES) and 5FOA-containing (YES+5FOA) media. Defects in heterochromatic gene silencing result in sensitivity to FOA, which counter-selects cells expressing *ura4*+. **D.**, **F.**, **H.** RT-qPCR analyses of *ura4*+ transcripts (mean±SD; n=3; normalized to *act1*+; relative to wt) produced from the different heterochromatic loci in cells of the indicated genetic background. Two-tailed Student’s *t*-tests were used to calculate *p*- values. Individual data points are represented by black circles.

Strikingly, however, silencing at the mating type locus was completely abolished in the absence of Caf1 and Mot2, similar to *clr4*Δ cells, as revealed by the lack of cell growth on 5FOA-containing medium and the marked accumulation of *ura4*+ transcripts (**Fig 1G, 1H**). These results were corroborated using strains carrying the *ade6*+ or *gfp*+ reporter genes instead of *ura4*+ (i.e. mat3M::*ade6*+ and mat3M::*gfp*+). Indeed, while pink/red colonies were observed for the wt strain on adenine-limiting medium, reflecting *ade6*+ silencing, *caf1*Δ and *mot2*Δ cells formed white colonies, like the *clr4*Δ mutant, indicative of derepressed *ade6*+ expression (**S1B Fig**). Steady state *gfp*+ mRNA levels were also strongly increased in these mutants, which correlated with the accumulation of the Gfp protein itself, supporting that heterochromatic transcripts are efficiently exported and translated in these different genetic backgrounds (**S1C, S1D Fig**). We also detected intermediate phenotypes for the *ccr4*Δ mutant, pointing to a partial contribution of this RNA deadenylase (**Figs 1G, 1H** and **S1B-D**). Moreover, endogenous matMc transcripts strongly accumulated in matP cells lacking Caf1 and Mot2, confirming the requirement for these factors in suppressing the expression of heterochromatic RNAs produced from the mating type locus (**S1E Fig**). We concluded from these experiments that the RNA deadenylase Caf1 and the E3 ubiquitin ligase Mot2 subunits of the Ccr4-Not complex are major regulators of heterochromatic gene silencing at the mating type locus.

Previous studies, including ours, showed that the Ccr4-Not complex tightly associates with the RNA-binding protein Mmi1 and its partner Erh1 to promote facultative heterochromatin assembly at meiotic genes and rDNA silencing [38-40,42-44]. We investigated whether these factors also contribute to gene silencing at the mating type locus and found that they were not required (**S1F, S1G Fig**). Hence, Ccr4-Not acts independently of Mmi1 and Erh1.

### The Ccr4-Not complex impacts heterochromatin assembly at the mating type locus

We next sought to determine whether defective silencing at the mating type locus in Ccr4-Not mutants is accompanied by an alteration in heterochromatin structural components. ChIP experiments revealed that the absence of Caf1 only modestly impacts H3K9me2 at mat3M::*ura4*+, while the *mot2*Δ mutant exhibited a pronounced decrease, albeit lower than that of *clr4*Δ cells (**Fig 2A**). We also assessed H3K9me3 levels and observed a further reduction in both *caf1*Δ and *mot2*Δ cells when compared to the wt strain (**Fig 2B**). Importantly, these defects were restricted to the mat3M locus, as H3K9me2/3 levels at pericentromeric dg repeats and subtelomeric *tlh1*+ sequences remained similar to those detected in wt cells, with the exception of a small H3K9me2 decrease at dg in the absence of Mot2 (**Fig 2A, 2B**). Total H3 levels were similar in all strains (**S2A Fig**), excluding an indirect effect linked to nucleosome instability. Since H3K9me2/3 constitutes a docking site for proteins of the HP1 family, we next analyzed the recruitment of the chromodomain-containing RITS subunit Chp1 involved in RNAi [45,46]. Following a similar trend, Chp1 occupancy was reduced specifically at mat3M::*ura4*+ in Ccr4-Not mutants (**Fig 2C**). Together, our results unveil a role for Caf1 and Mot2 in heterochromatin assembly, which correlates with their requirement for gene silencing at the mating type locus (**Fig 1G, 1H**). They further suggest that both factors operate differentially, Mot2 being more critical for heterochromatin integrity.

**Fig 2.**
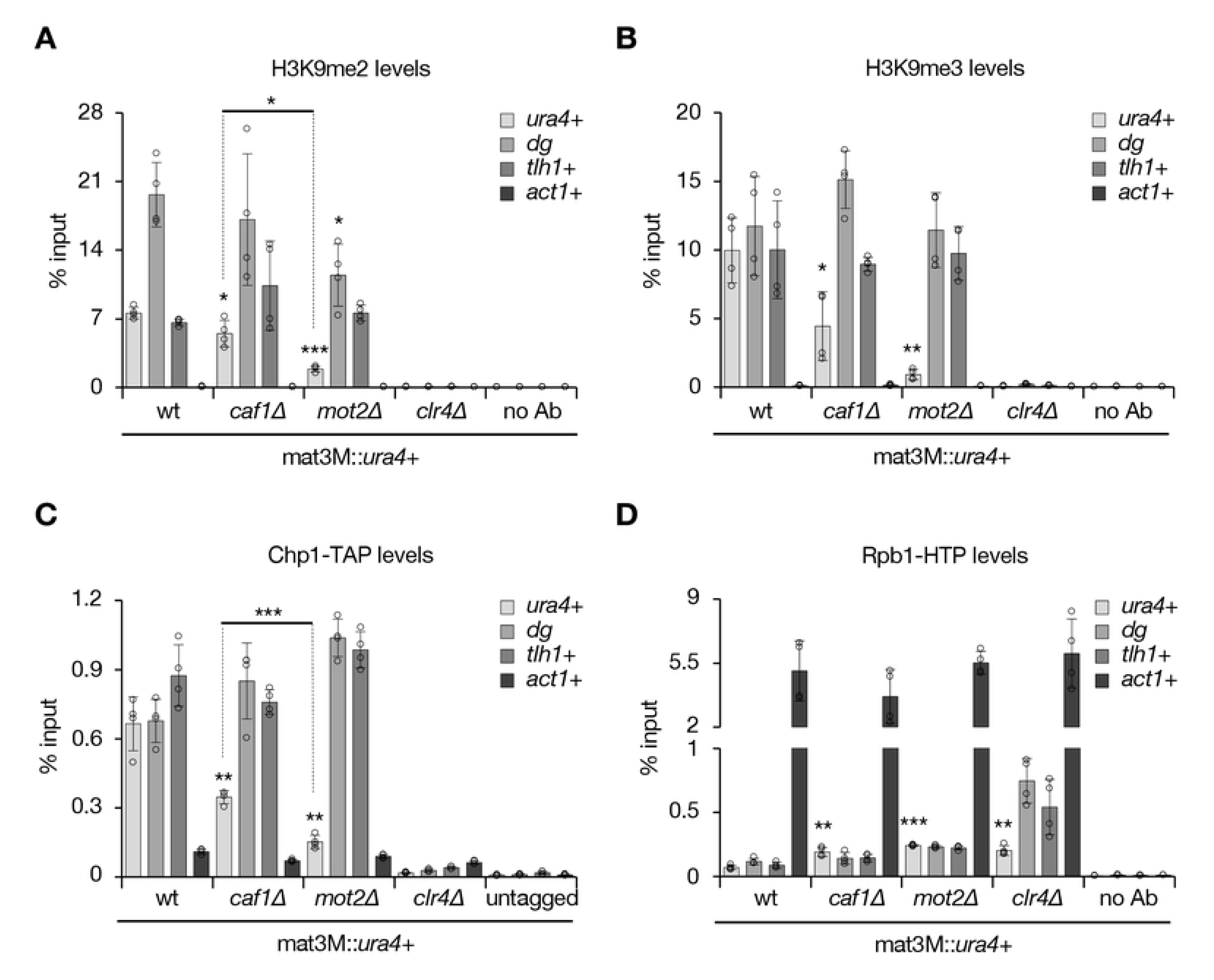
The Ccr4-Not complex impacts heterochromatin assembly at the mating type locus. **A.-D.** ChIP-qPCR analyses (% input; mean±SD; n=4) of the indicated histone modifications and tagged proteins in mat3M::*ura4*+ strains. Shown are the enrichments of *ura4*+, *dg* pericentromeric repeats, the *tlh1*+ subtelomeric locus and *act1*+ upon immunoprecipitation with H3K9me2 (**A.**), H3K9me3 (**B.**) antibodies or rabbit IgG (**C.**, **D.**). Immunoprecipitations without antibodies (no Ab) or from untagged strains were performed to determine background levels. In **D.**, “no Ab” values represent the mean of two independent analyses (n=2). Two-tailed Student’s *t*-tests were used to calculate *p*-values. Individual data points are represented by black circles.

Clr4-mediated heterochromatin assembly at centromeres and telomeres is essential to limit RNAPII accessibility and hence trigger efficient TGS, whereas its impact on the transcription machinery at the mating type locus is moderate, reflecting a predominance of C/*cis*-PTGS [30]. In agreement with these notions, we found that RNAPII strongly accumulated at dg and *tlh1*+ sequences but only modestly at mat3M::*ura4*+ in the absence of Clr4 (**Fig 2D**). In *caf1*Δ and *mot2*Δ mutants, RNAPII levels increased similarly at mat3M but only marginally at pericentromeric or subtelomeric loci when compared to *clr4*Δ cells (**Fig 2D**), consistent with the relative impact of Ccr4-Not in silencing/heterochromatin formation at these different regions. We also assessed whether Caf1 and Mot2 themselves are recruited to the mating type locus but failed to detect a significant enrichment, as opposed to the RITS component Chp1 (**S2B Fig**). Thus, Caf1 and Mot2 are not stably recruited to heterochromatin, which might indicate that their association is too transient, as suggested by the weak physical interaction between the Ccr4 subunit and Chp1 [39].

### The Ccr4-Not subunits Caf1 and Mot2 regulate heterochromatin spreading

The transition from H3K9me2 to H3K9me3 contributes to heterochromatin spreading from nucleation sites [25,26,29]. Our findings that *caf1*Δ and *mot2*Δ cells predominantly affect H3K9me3 at mat3M::*ura4*+ prompted us to examine their potential impact on heterochromatin spreading. Intriguingly, ChIP experiments revealed a gradual decrease of H3K9me3 in Ccr4-Not mutants relative to the wt strain, from the cenH nucleation center up to inverted repeats at the right border (IR-R) of the mating type locus (**Fig 3A**). Consistent with the above results, the effect was also more prominent in the absence of Mot2.

**Fig 3.**
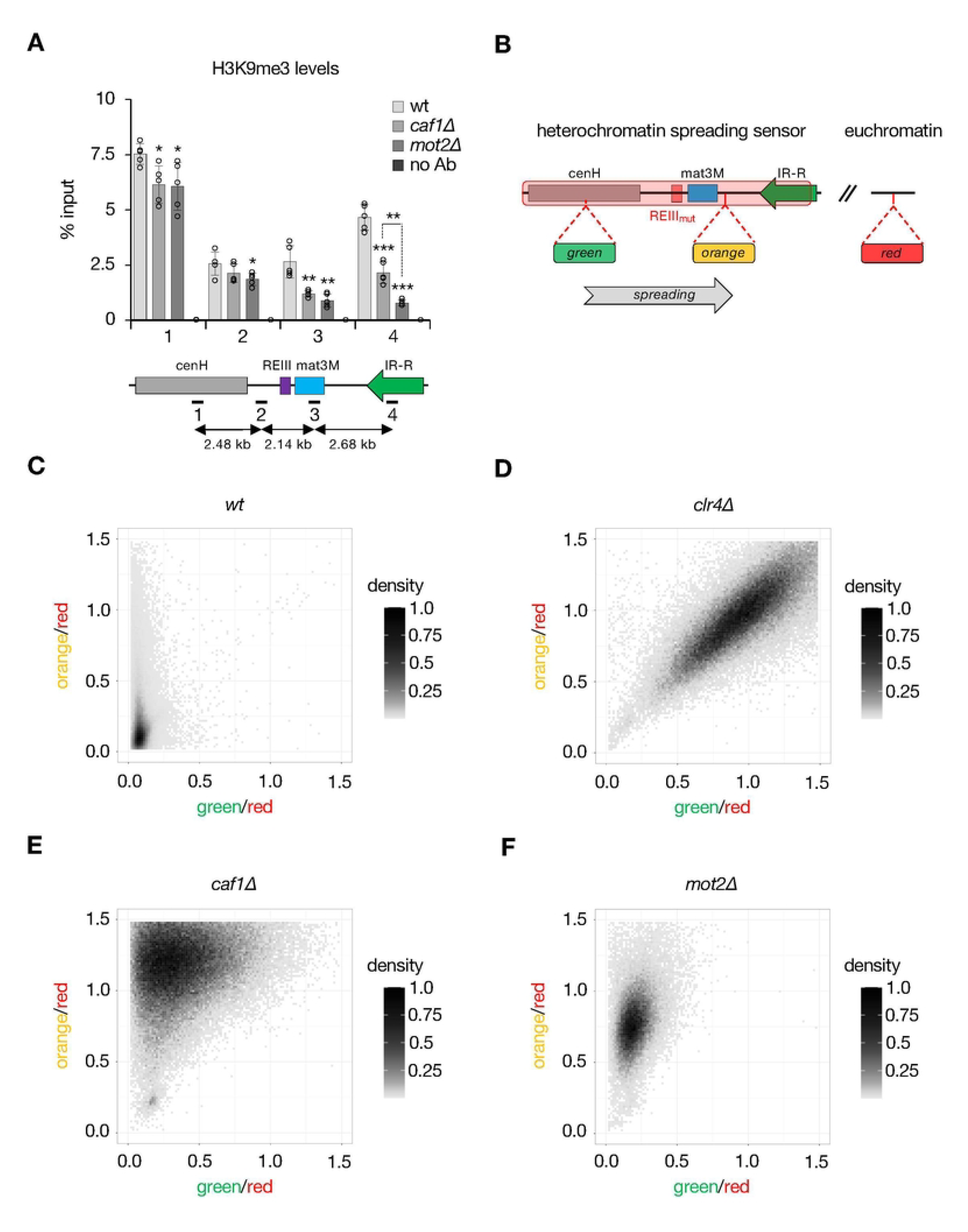
The Ccr4-Not subunits Caf1 and Mot2 regulate heterochromatin spreading. **A**. H3K9me3 ChIP-qPCR analyses (% input; mean±SD; n=5) in cells of the indicated genetic backgrounds. Numbers correspond to the different primer pairs used in qPCR reactions and whose localization is indicated on the scheme below the graph. Immunoprecipitations without antibodies (no Ab) were performed to determine background levels. Two-tailed Student’s *t*- tests were used to calculate *p*-values. Individual data points are represented by black circles. **B**. Scheme depicting the heterochromatin spreading sensor with the relative positions of the “green”, “orange” and “red” reporters. **C.-F.** Two-dimensional-density squarebin plots showing the red-normalized green and orange fluorescence for wt (**C.**), *clr4Δ* (**D.**), *caf1Δ* (**E.**) and *mot2*Δ (**F.**) REIII_mut_ cells grown at 32°C. A density bar represents the fraction of the most dense bin. One representative isolate is shown for each background.

To substantiate these findings, we used our previously described fluorescent reporter-based, single-cell spreading sensor assay [47,48], whereby three different fluorescent protein-coding genes allow quantitative measurements of gene expression at nucleation-proximal (“green”) and -distal (“orange”) heterochromatin sites, as well as at a euchromatic locus (“red”) for signal normalization purposes (**Fig 3B**). Because heterochromatin at the mating type locus can nucleate from two distinct regulatory DNA elements (i.e. cenH and REIII), we analyzed strains in which the Atf1/Pcr1-binding sites within REIII are mutated (REIII_mut_), thereby allowing to record heterochromatin spreading (“orange”) from the sole cenH nucleation center (“green”) (**Fig 3B**).

Flow cytometry analyses revealed that wt REIII_mut_ cells successfully nucleated heterochromatin (“green”^OFF^), which efficiently spread to the distal reporter (“orange”^OFF^), as indicated by the strong enrichment of cell populations in the bottom left part of the 2D density squarebin plot (**Fig 3C**). A minor fraction of cells displayed some “orange” signal though, reflecting a certain degree of stochasticity in the spreading process [47,49]. Upon complete loss of heterochromatin and silencing (i.e. in *clr4*Δ cells), populations concentrated along the diagonal, in the upper right part of the plot, consistent with both reporters being fully derepressed (“green”^ON^ and “orange”^ON^) (**Fig 3D**). Remarkably, Ccr4-Not mutants exhibited radically different patterns, with a strong bias for “orange” versus “green” fluorescence overall (**Figs 3E, 3F** and **S3A-D**). In the absence of Caf1, the majority of cells distributed within a broad range of high “orange” and intermediate “green” signals (**Figs 3E** and **S3A-B**). This indicates that the deadenylase strongly suppresses the expression of the nucleation-distal reporter while only partially contributing to the silencing of the proximal locus. Of note, we reproducibly observed a subpopulation in the bottom left part of the plot, indicating that some cells maintained both reporters in the repressed state. Cells lacking Mot2 showed instead a narrow, vertical distribution of fluorescence, covering a large range of high “orange” intensities associated with low “green” signals (**Figs 3F** and **S3C-D**). Such pattern is reminiscent to what previously observed for *bona fide* regulators of heterochromatin spreading [49]. Consistent with these results, cenH proximal-transcripts were only partially increased in Ccr4-Not mutants when compared to *clr4*Δ cells (**S3E Fig**), in striking contrast to distal matMc RNAs (**S1E Fig**).

Because the pericentromeric and subtelomeric reporter strains used in this study (i.e. otr1R::*ura4*+ and tel2L::*ura4*+; **Fig 1C-1F**) carry the *ura4*+ gene in proximity to nucleation sites (i.e. dg and *tlh1*+ sequences, respectively), we next investigated heterochromatin propagation at these regions by ChIP. While H3K9me3 distribution remained similar to wt cells at pericentromeres (**S4A Fig**), *caf1*Δ and *mot2*Δ mutants exhibited lower levels away from the subtelomeric *tlh1*+ gene (**S4B Fig**), consistent with former observations for H3K9me2 [34, 39]. Expression analyses further revealed that, relative to Clr4, Caf1 had a more prominent role in the suppression of distal subtelomeric transcripts (**S4C Fig**). The contribution of Mot2 followed a similar tendency, although the most distal genes were instead downregulated in its absence, likely due to additional effects impairing expression of these loci (**S4C Fig**).

Overall, our data establish critical, yet different roles for Caf1 and Mot2 in the propagation of H3K9me3 and the repression of nucleation-distal transcripts, pointing to key contributions of Ccr4-Not to heterochromatin spreading at the mating type locus and subtelomeres.

### Importance of Caf1 and Mot2 catalytic activities in gene silencing and heterochromatin spreading

To determine the mechanisms by which Caf1 and Mot2 mediate heterochromatic gene silencing and spreading, we assessed the impact of mutants defective for their RNA deadenylation and E3 ubiquitin ligase activities, which are carried by Ribonuclease H superfamily and RING-type domains respectively (**Fig 4A**).

**Fig 4.**
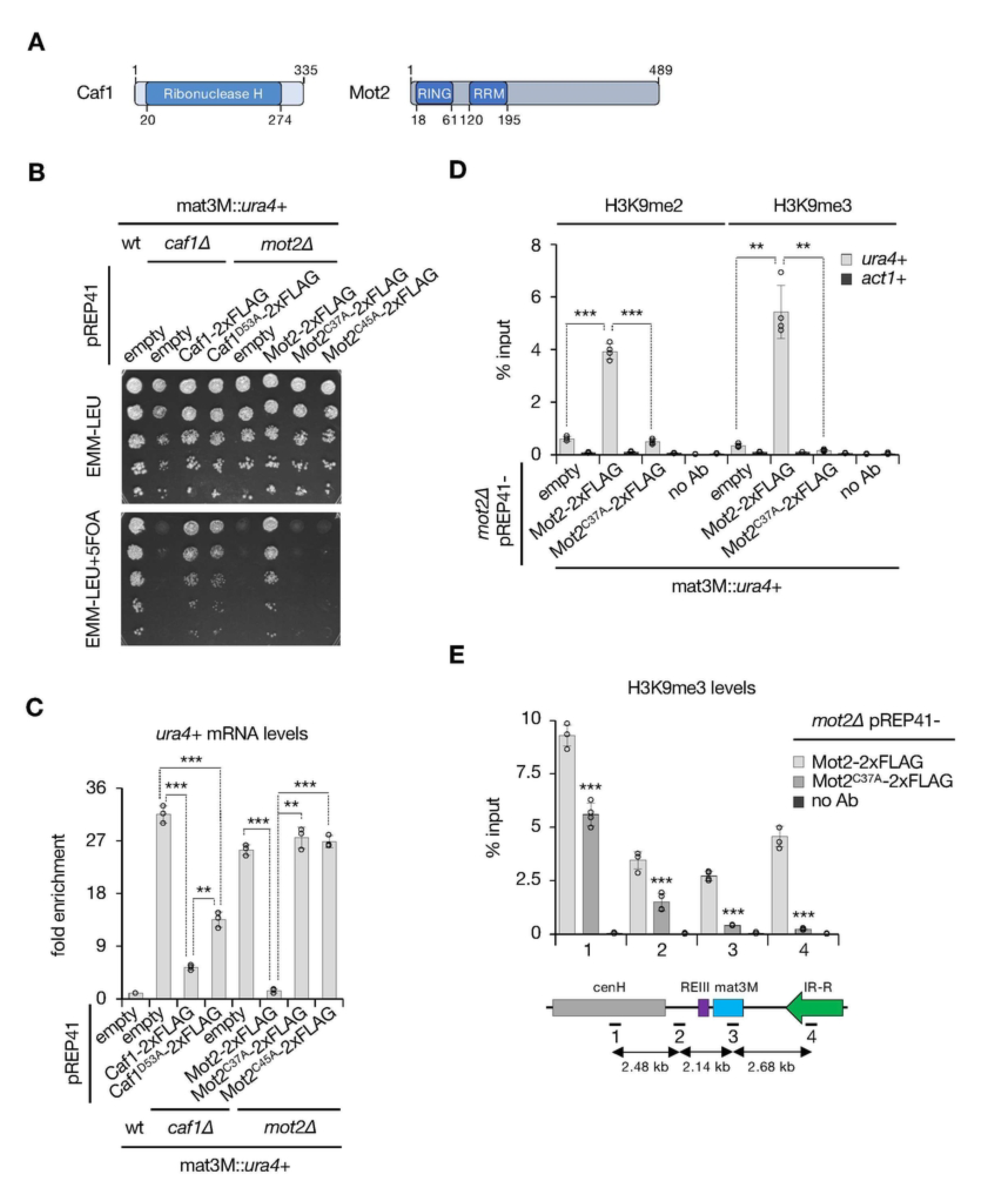
Importance of Caf1 and Mot2 catalytic activities in gene silencing and heterochromatin spreading. **A.** Domain organization of Caf1 and Mot2 proteins. **B.** 5-fold serial diluted silencing assay using the *ura4*+ reporter gene inserted at the mating type locus (mat3M::*ura4*+). Cells of the indicated genotypes were plated on both non-selective (EMM- LEU) and 5FOA-containing (EMM-LEU+5FOA) media. **C.** RT-qPCR analyses of *ura4*+ transcripts (mean±SD; n=3; normalized to *act1*+; relative to wt pREP41) in cells of the indicated genetic backgrounds. **D.**, **E.** H3K9me2 and H3K9me3 ChIP-qPCR analyses (% input; mean±SD; n=4 or 3) in cells of the indicated genetic backgrounds. Immunoprecipitations without antibodies (no Ab) were performed to determine background levels. In **E.**, numbers correspond to the different primer pairs used in qPCR reactions and whose localization is indicated on the scheme below the graph. **C.**-**E.** Two-tailed Student’s *t*-tests were used to calculate *p*-values. Individual data points are represented by black circles.

We first expressed a plasmid-borne, 2xFLAG-tagged version of Caf1^D53A^ mutant, previously shown to impair RNA deadenylation [50], in *caf1*Δ mat3M::*ura4*+ cells (**S5A Fig**). Although we observed only a mild growth defect in the presence of 5FOA (**Fig 4B**), *ura4*+ mRNA levels were significantly increased, yet not as pronounced as *caf1*+ deletion (**Fig 4C**). This likely reflects the contribution of the second deadenylase Ccr4 (**Figs 1H** and **S1B-D**) [34], which is physically tethered to the complex by Caf1 [51] and hence does not exert its function in *caf1*Δ cells. Of note, a plasmid-borne, 2xFLAG-tagged wild type Caf1, did not allow to fully restore growth on 5FOA nor low levels of *ura4*+ transcripts in *caf1*Δ cells (**Fig 4B, 4C**), indicating that ectopic expression of Caf1 also partially inhibits silencing. Overall, these results indicate that the RNA deadenylation activity of Caf1 partially contributes to heterochromatic gene silencing.

Next, we assessed the requirement for Mot2 catalytic activity following the same strategy. Expression of a plasmid-borne Mot2^RINGΔ^ mutant in otherwise *mot2*Δ cells completely abolished *ura4*+ silencing, as determined by the lack of growth in the presence of 5FOA and RT-qPCR analyses (**S5B, S5C Fig**). We further studied mutants in which key cysteine residues in the RING domain were substituted by alanine (Mot2^C37A^, Mot2^C45A^) [42] and similarly observed the absence of growth on 5FOA-containing medium as well as a strong accumulation of *ura4*+ mRNAs, akin to the *mot2*Δ strain (**Fig 4B, 4C**). Importantly, these phenotypes did not result from a lowered expression of the mutant proteins (**S5D Fig**). Since the absence of Mot2 strongly impairs heterochromatin assembly at the mating type locus (**Fig 2A, 2B**), we also investigated the impact of Mot2^C37A^ on H3K9me at this region. ChIP analyses revealed a marked decrease in both H3K9me2 and H3K9me3 levels at mat3M::*ura4*+, similar to *mot2*Δ cells (**Fig 4D**). We further assessed the distribution of H3K9me3 from the cenH nucleation center to inverted repeats and reproducibly observed a progressive decrease (**Fig 4E**), strongly suggesting a defect in the spreading process. Hence, we concluded that the E3 ubiquitin ligase activity of Mot2 has a major role in gene silencing, heterochromatin assembly and spreading.

### The anti-silencing factor Epe1 opposes Ccr4-Not in gene silencing and heterochromatin spreading

Previous studies showed that deletion of the JmjC domain-containing protein Epe1 suppresses silencing defects observed in mutants of the heterochromatin machinery [7,8,52-55]. To determine whether Epe1 also opposes the function of Ccr4-Not, we constructed *caf1*Δ *epe1*Δ and *mot2*Δ *epe1*Δ double mutants and assessed their ability to restore heterochromatic gene silencing at the mating type locus. Interestingly, silencing defects in the absence of Caf1 and Mot2 were completely suppressed upon *epe1*+ deletion (**Fig 5A**). This was in marked contrast to *clr4*Δ cells, consistent with the notion that H3K9me is required for Epe1 recruitment. RT-qPCR assays and Northern blotting further confirmed that the absence of Epe1 restricts the accumulation of *ura4*+ mRNAs in *caf1*Δ and *mot2*Δ, but not *clr4*Δ mutants (**Figs 5B** and **S6A**). Likewise, defects in H3K9me2 and H3K9me3 in cells lacking Mot2 were alleviated upon loss of Epe1 (**Fig 5C**), further indicating that the latter mediates the heterochromatin assembly defects. These results also suggest that *caf1*Δ and *mot2*Δ cells maintain sufficient levels of H3K9me to ensure Epe1 recruitment. Indeed, ChIP experiments revealed that the protein was similarly enriched at mat3M::*ura4*+ when compared to the wt strain (**Fig 5D**), and Western blot analyses further excluded an alteration in Epe1 protein levels in the mutants (**S6B Fig**).

**Fig 5.**
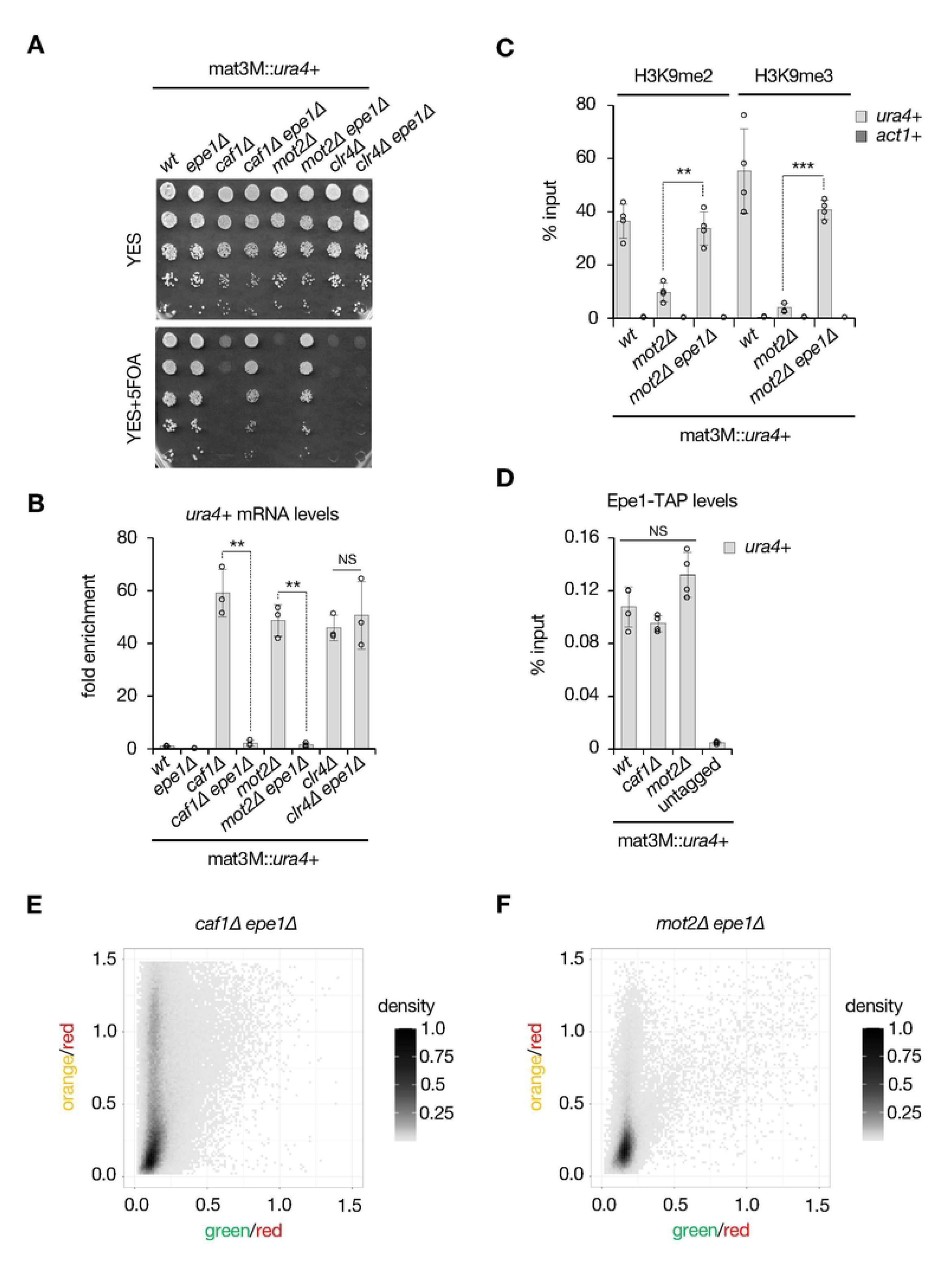
The anti-silencing factor Epe1 opposes Ccr4-Not in gene silencing and heterochromatin spreading. **A.** 5-fold serial diluted silencing assays using the *ura4*+ reporter gene inserted at the mating type locus (mat3M::*ura4*+). Cells of the indicated genotypes were plated on both non-selective (YES) and 5FOA-containing (YES+5FOA) media. **B.** RT-qPCR analyses of *ura4*+ transcripts (mean±SD; n=3; normalized to *act1*+; relative to wt) in cells of the indicated genetic backgrounds. **C.** H3K9me2 and H3K9me3 ChIP-qPCR analyses (% input; mean±SD; n=4) in cells of the indicated genetic backgrounds. **D.** Epe1-TAP ChIP-qPCR analyses (% input; mean±SD; n=4) in cells of the indicated genetic backgrounds. An untagged strain was used as negative control. **B.-D.** Two-tailed Student’s *t*-tests were used to calculate *p*- values. Individual data points are represented by black circles. **E.**, **F.** Two-dimensional-density squarebin plots showing the red-normalized green and orange fluorescence for *caf1Δ epe1Δ* (**E.**) and *mot2*Δ *epe1Δ* (**F.**) REIII_mut_ cells grown at 32°C. A density bar represents the fraction of the most dense bin. One representative isolate is shown for each background.

To determine the contribution of Epe1 to the heterochromatin spreading defects observed in Ccr4-Not mutants, we next performed flow cytometry analyses in the *caf1*Δ *epe1*Δ and *mot2*Δ *epe1*Δ strains as described above. In both cases, cell populations with high spreading marker (“orange”) fluorescence were strongly diminished compared to single mutants, yet not fully to wt levels (**Fig 5E, 5F**). In particular, *caf1*Δ *epe1*Δ populations still distributed over a broad range of “orange” signals, while “green” fluorescence was comparable to what observed in the wt. *mot2*Δ *epe1*Δ cells also displayed a pronounced reduction in “orange” and maintained low levels of “green”. We concluded from these experiments that the loss of Epe1 strongly limits the extent and penetrance of heterochromatin spreading defects in Ccr4-Not mutants, akin to its effects on gene repression and heterochromatin formation.

To get additional insights into the mechanism by which it antagonizes Ccr4-Not, we subsequently generated an Epe1 mutant in the JmjC domain (i.e. Epe1^H297A^) that was inserted at its endogenous locus and expressed from its own promoter (**S7A Fig**). Combining this allele to the deletion of *caf1*+ or *mot2*+ fully restored cell growth in the presence of 5FOA and suppressed the accumulation of *ura4*+ mRNAs produced from the mat3M locus (**S7B, S7C Fig**). We verified by Western blotting that the mutant protein was similarly expressed to the wt version in the different genetic backgrounds (**S7D Fig**). Consistent with former work [56], ChIP experiments further indicated that Epe1^H297A^ fails to be efficiently recruited to heterochromatin (i.e. mat3M::*ura4*+) (**S7E Fig**), providing a rationale for the suppression of silencing defects in *caf1*Δ and *mot2*Δ cells. Thus, the loss of heterochromatic gene silencing in mutants of the Ccr4-Not complex strictly depends on the integrity of the Epe1 JmjC domain.

## Discussion

In this study, we demonstrate that the Caf1 and Mot2 subunits of Ccr4-Not are two major catalytic effectors of heterochromatic gene silencing and spreading. Our findings not only go beyond former observations by establishing an essential role for the complex in silencing at the mating type locus but also add another layer to its regulatory control, i.e. the implication of its catalytic components in heterochromatin propagation.

The nuclease module of the Ccr4-Not complex comprises the Caf1 and Ccr4 deadenylases, the former anchoring the latter to the Not1 scaffolding subunit [51]. Consistent with this and previous analyses implicating the nuclease activity of both enzymes in RNA degradation and heterochromatic gene silencing [34,50], we found that i) catalytically inactive Caf1 elicits weaker defects than its deletion and, ii) Ccr4 partially contributes to the gene silencing activity. Caf1 and Ccr4 may thus act in concert onto polyadenylated, heterochromatic RNAs, although it remains unclear whether they target common or individual transcripts. Both enzymes may further promote the recruitment of additional RNA degradation factors important for silencing activities, as previously suggested [34]. The fact that the heterologous mat3M::*gfp*+ reporter is derepressed in the absence of Caf1 also makes it unlikely that defined factor(s) tether the complex to heterochromatic transcripts in a sequence-dependent manner. It is however possible that poly(A)-binding proteins and/or the Mot2 subunit itself, which carries an RNA Recognition Motif, may recruit Ccr4-Not to promote deadenylation [57-59]. Alternatively, the complex may directly associate with the transcription machinery, as illustrated by its interaction in budding yeast with RNAPII and the histone chaperone Spt6 [60,61], both of which also contribute to heterochromatic gene silencing in fission yeast [62-65]. Regardless the precise modalities, the recruitment of Ccr4-Not deadenylases is essential to clear RNAs from chromatin, thereby maintaining heterochromatin integrity (this study; [34]).

Our work also establishes a potent, catalytic role for the E3 ubiquitin ligase Mot2 in gene silencing and heterochromatin assembly. A parsimonious model predicts that Mot2 may target an anti-silencing factor for ubiquitinylation-dependent proteasomal degradation, as previously described for the E3 ligase Ddb1 that promotes Epe1 turnover and restricts its accumulation within heterochromatin domains [52]. However, this factor is unlikely to be such a substrate given its similar abundance and occupancy in wt and *mot2*Δ cells. Mot2 could instead target other functionally-related factors, including the bromodomain-containing protein Bdf2 and the histone acetyltransferase Mst2 [11,66,67], thereby shielding heterochromatin from invading anti-silencing activities. Other scenarios include the modification of RNAPII itself, perhaps to stimulate elongation on heterochromatin templates, akin to situations where RNAPII transiently pauses or encounters transcriptional blocks [60,68-70]. Persistence of the transcription machinery, as suggested by our analyses, and/or the accumulation of transcripts in the vicinity of the DNA template may in turn alter heterochromatin structure [34]. Interestingly, Ccr4-Not components, including Mot2 and Caf1, were recently proposed to regulate transcriptional efficiency at heterochromatic loci by limiting the levels of RNAPII-associated transcripts [41]. The causal relationship between this process and silencing efficiency (i.e. steady state RNA levels) remains however opaque.

Another critical aspect of our findings is the requirement for both Caf1 and Mot2 in heterochromatin spreading. This is supported by several lines of evidence in mutant cells: i) H3K9me3 is further reduced than H3K9me2 at the mat3M locus, consistent with a defect in the transition between these two heterochromatin states, ii) propagation of H3K9me3 from nucleation centers is impaired at the mating type locus and subtelomeres, iii) nucleation-distal reporters/genes are further derepressed than proximal loci, and iv) the absence of Epe1, which antagonizes heterochromatin spreading, suppresses the observed silencing and spreading defects. Further supporting a role for Ccr4-Not in heterochromatin spreading, the Ccr4 and Rcd1 subunits were also recently identified in our screen for modulators of spreading at the mating type locus [49]. From a mechanistic perspective, however, the exact role of the complex still needs to be elucidated. It is possible that elimination of RNAPII/chromatin-associated transcripts by Ccr4 and Caf1 facilitates the transition from transcriptionally-permissive H3K9me2 to H3K9me3 by Clr4 [29,34,41], thereby ensuring efficient spreading. Their minor contribution to RNA degradation at nucleation centers (i.e. cenH and *tlh1*+) underlies functional redundancy with RNAi [34], yet deadenylases may take over at distal loci due to the lack of siRNA templates. As for Mot2, the enzyme could, beyond the scenarios evoked above, directly control the activity of spreading regulators [49,55], possibly through non-proteolytic ubiquitinylation. Future studies aimed at identifying the substrate of Mot2 will be instrumental to understand the molecular basis of its involvement in heterochromatin spreading.

Despite reduced H3K9me2/3 levels, *caf1*Δ and *mot2*Δ cells maintain Epe1 occupancy at the mat3M locus, suggesting that the balance between pro- and anti-silencing activities is altered. Perhaps Epe1 facilitates heterochromatin transcription and/or histone turnover in these genetic contexts [8,53,71], thereby accounting for the observed mutant phenotypes. Our data showing that the protein opposes Ccr4-Not in a JmjC domain-dependent manner might also indicate that failure to antagonize H3K9me allows restoring silencing and spreading activities. However, the JmjC domain of Epe1 does not support H3K9 demethylase activity *in vitro* but is instead crucial for a robust association with Swi6/HP1 *in vivo* [56,72]. It is therefore conceivable that the disrupted interaction between Epe1^H297A^ and Swi6 is the main reason why silencing is re-established in Ccr4-Not mutants. Indeed, Epe1^H297A^ is not properly recruited to HP1-coated heterochromatin (this study; [56]), which may restrict anti-silencing activities and hence ensure efficient H3K9me3 maintenance and spreading.

Heterochromatin domains are spatially segregated at the nuclear periphery within distinct sub-compartments [73-75]. The nuclear envelope protein Amo1 was shown to anchor the mating type locus and to stably associate with the Ccr4-Not complex [73], raising the possibility that the latter may also regulate perinuclear sequestration of this heterochromatin domain. Whether additional peripheral proteins may similarly interact with Ccr4-Not to co-regulate telomere positioning remains an open question. Beyond these considerations, Amo1 cooperates with the rixosome and the histone chaperone FACT to mediate RNA degradation and suppress histone turnover for proper silencing and epigenetic inheritance of heterochromatin [73,76,77]. Intriguingly, Ccr4 was identified in a screen for factors involved in heterochromatin inheritance and Caf1 was found to act in parallel to the rixosome for degradation of heterochromatic RNAs [76], pointing to a tight relationship between these machineries. Whether Ccr4-Not also partakes in histone turnover and epigenetic inheritance of heterochromatin are fascinating possibilities requiring further investigation.

In conclusion, we demonstrate a fundamental role for the highly-conserved Ccr4-Not complex in gene silencing, heterochromatin assembly and spreading, which implicates its Caf1 and Mot2 subunits. Our findings open new perspectives to dissect the deadenylation- and ubiquitinylation-dependent mechanisms involved and assess the functional and biological relevance of such activities in disease-related models. The cooperation between human CCR4-NOT and the H3K9me3-functionally linked HUSH complex in HIV repression provides a meaningful example in this respect [78,79].

## Materials and Methods

### Strains, media and plasmids

The *S. pombe* strains used in this study are listed in **S1 Table**. Strains were generated by transformation following a lithium acetate-based method or by random spore analysis (mating and sporulation on Malt Extract) using complete medium (YE Broth, Formedium, #PMC0105) supplemented with appropriate antibiotics. Experiments were performed in 1X YE supplemented with 150 mg/L of adenine (Sigma, #A2786), L-histidine (Sigma, #H8000), L-lysine (Sigma, #L5501), L-leucine (Sigma, #L8000) and uracile (Sigma, #U750) (YES), while plasmids-containing strains were grown in 1X EMM-LEU-URA minimal medium (EMM-LEU-URA Broth, Formedium, #PMD0810) supplemented with uracil (Sigma, #U750) (EMM-LEU). For silencing assays, cells were grown until mid-log phase and plated on both non-selective (YES or EMM-LEU) and selective (YES+5FOA, EMM-LEU+5FOA or YES low ADE) media at an initial OD = 0.2 – 0.3 followed by 5-fold serial dilutions. Pictures were taken after 3 to 4 days of growth at 30°C.

The plasmids used for gene cloning/editing are listed in **S2 Table**. To construct strains expressing tagged versions of wt or mutant Epe1 from its genomic locus, we first deleted the ORF with a cassette containing the hygromycin resistance marker (hph^R^MX) fused to the herpes simplex virus thymidine kinase-encoding gene (HSV-TK) from the pFA6a-HyTkAX vector (Addgene plasmid # 73898; http://n2t.net/addgene:73898; RRID:Addgene_73898) [80]. The resulting hph^R^ and TK-expressing *epe1*Δ strain was then transformed with a PCR product of interest (i.e. Epe1-TAP or Epe1^H297A^-TAP) carrying homology with the promoter and terminator regions. Positive integrants were selected in the presence of 5-fluoro-2’- desoxyuridine, which counter-selects cells expressing TK, and became again hygromycin-sensitive due to cassette pop-out.

### RNA extraction

Total RNAs were extracted from 4 mL of yeast cells at OD=0.8-1.0. Following centrifugation, cells were washed in water and frozen in liquid nitrogen. Cell pellets were resuspended in 1 volume of TES buffer (10 mM Tris-HCl pH7.5, 5mM EDTA, 1% SDS) and 1 volume of acid phenol solution pH4.3 (Sigma, #P4682), and incubated for 1 hour at 65°C in a thermomixer with shaking. After centrifugation, the aqueous phase was recovered and 1 volume of chloroform (ThermoFisher Scientific, #383760010) was added. Samples were vortexed and centrifugated followed by ethanol precipitation in the presence of 200 mM of lithium acetate. Pellets were resuspended in water and treated with DNAse (Ambion, #AM1906). RNA concentrations were measured with a Nanodrop.

### Reverse Transcription and real-time PCR (RT-qPCR)

2 µg of DNAse-treated RNAs were denatured at 65°C for 5 minutes in the presence of strand-specific primers or a mix of random hexamers (ThermoFisher Scientific, #SO142) and oligodT. Reactions were carried out with 100 units Maxima Reverse Transcriptase (ThermoFisher Scientific, #EP0743) at 50°C for 30 minutes. The enzyme was then denatured at 85°C for 5 minutes, and reactions diluted to 1:10 ratio. Each experiment included negative controls without Reverse Transcriptase. Samples were quantified by qPCR with SYBR Green Master Mix and a LightCycler LC480 apparatus (Roche). Measurements were statistically compared using two-tailed t-tests with the following p-value cut-offs for significance: 0.05>*>0.01; 0.01>**>0.001; ***<0.001. Oligonucleotides used in qPCR reactions are listed in **S3 Table**.

### Chromatin Immunoprecipitation and real-time PCR (ChIP-qPCR)

40 to 50 mL yeast cultures were grown in appropriate medium until OD=0.8-1.0. Crosslinking was performed by adding 1% formaldehyde (Sigma, #F8775) for 20 min at 30°C on the shaker and the reaction was quenched with 250 mM glycine (Sigma, #G7126) for 5 min at room temperature. Cells were washed with 1X ice-cold PBS, harvested by centrifugation and frozen in liquid nitrogen. Pellets were resuspended in 500 µL 150 mM FA buffer (50 mM Hepes-KOH pH7.5, 150 mM NaCl, 1 mM EDTA, 1% Triton X-100, 0.1% Na deoxycholate) in 2 mL screw-cap tubes. 1 mL of ice-cold, acid-washed glass beads (Sigma, #G8772) was added, and cells were lysed by 6 cycles of 40 sec at 6000 rpm using a FastPrep-24 5G apparatus (MP). Following centrifugation, pellets were resuspended in 150 mM FA buffer and sonicated for 6 cycles of 30 sec at 40% amplitude using a tip probe VibraCell sonifier (Bioblock Scientific). Chromatin extracts were then cleared by centrifugation for 15 min at 14000 rpm at 4°C. 100 µL was set aside as the input control and 200 to 400 µL aliquots were typically used for immunoprecipitations. 1 to 2 µL antibodies against total H3 (Abcam, ab1791), H3K9me2 (Abcam, ab1220) and H3K9me3 (Abcam, ab8898) were added to the lysates and samples were incubated at 4°C on a wheel for 2 hours or over-night. 4 µL pre-washed ProteinA or ProteinG Dynabeads (Invitrogen, #10001D or #10003D) were next added and samples were incubated for additional 2 hours at 4°C. For immunoprecipitation of TAP or HTP-tagged proteins (Chp1-TAP, Rpb1-HTP and Epe1-TAP), 4 µL of pre-washed rabbit IgG-conjugated M-270 Epoxy Dynabeads (Invitrogen, #14311D) were added and lysates were incubated on a wheel for 1 hour at 4°C. Magnetic beads were then washed at room temperature twice with 150 mM FA buffer, twice with 500 mM FA buffer, once with wash buffer (10 mM Tris-HCl pH8, 0.25 M LiCl, 1mM EDTA, 0.5% NP-40, 0,5% Na deoxycholate) and once with TE buffer (10 mM Tris-HCl pH8, 1 mM EDTA). DNA was then eluted with 100 µL ChIP Elution buffer (50 mM Tris pH7.5, 10 mM EDTA, 1% SDS) for 20 min at 80°C in a Thermomixer set to 1400 rpm. 20 µg proteinase K (Euromedex, #09-911) were added to the input and IP samples for 30 min at 37°C prior to over-night decrosslinking at 65°C. The following day, 1 µL RNAseA/T1 mix (Thermo Scientific, #EN0551) was added for 30 min at 37°C and DNA was purified using NucleoSpin Gel and PCR Clean-Up columns (Macherey-Nagel, #740609.250) according to the manufacturer’s instructions. Samples were quantified by qPCR with SYBR Green Master Mix and a LightCycler LC480 apparatus (Roche). Measurements were statistically compared using two-tailed t-tests with the following p-value cut-offs for significance: 0.05>*>0.01; 0.01>**>0.001; ***<0.001. Oligonucleotides used in qPCR reactions are listed in **S3 Table**.

### Total protein analyses

Total proteins were extracted from cell pellets corresponding to 2 to 5 ODs, as described in [42]. Cell lysis was performed on ice using 0.3M NaOH and 1% beta-mercaptoethanol prior to protein precipitation with trichloroacetic acid (TCA) (7% final). Following full speed centrifugation, pellets were resuspended in HU loading buffer and heat-denaturated at 70°C. Soluble fractions were recovered, and samples were analyzed by standard immunoblotting procedures using 1:3000 peroxydase-conjugated antiperoxydase (PAP, to detect protein-A-tagged proteins) (Sigma, #P1291, RRID:AB_1079562), 1:3000 monoclonal anti-FLAG antibody (Sigma, #F3165, RRID:AB_259529), 1:3000 anti-CDC2 antibody (Abcam, #ab5467, RRID:AB_2074778), 1:1000 anti-GFP antibody (Roche, # 11814460001, RRID: AB_390913) and 1:5000 goat anti-mouse IgG-HRP (Santa Cruz Biotechnology, #sc-2005, RRID:AB_631736). Detection was done with SuperSignal West Pico Chemiluminescent Substrate (ThermoFisher Scientific, #34080), ECL Select reagent (GE Healthcare, #RPN2235), and a Vilber Lourmat Fusion Fx7 imager or a ChemiDoc Touch Imaging System (BIORAD).

### Northern blotting

10 µg RNAs were separated on a 1.2% agarose gel and transferred overnight by capillarity on a nylon membrane (GE Healthcare, #RPN203B). RNAs were then UV-crosslinked to the membrane using a Stratalinker apparatus. A radiolabeled PCR probe was prepared by random priming (GE Healthcare, Megaprime kit) using *α*-P^32^ dCTP and incubated with the membrane in commercial buffer (Ambion, UltraHyb). Following washes, the membrane was exposed for 24 hours and revelation was performed using a Typhoon phosphoimager. Oligonucleotides used to generate the probe are listed in **S3 Table**.

### Flow cytometry data collection and normalization for validation

Cells were struck out in YES, grown in liquid YES overnight to saturation and then diluted 1:15 prior to further growth in liquid YES up to mid log phase. Flow analysis was performed on a LSR Fortessa X50 (BD Biosciences) and color compensation, analysis and plotting were performed as described previously [49].

## Data Availability

All the raw data allowing to generate the final results presented here are available in supporting information as a separate excel file “Challal D 2023 Raw Data”.

The R script and primary fcs files used to establish flow cytometry data have been deposited in Zenodo (doi: 10.5281/zenodo.7611852).

## Acknowledgments

We thank Marc Bühler, Alain Jacquier, Jean-Paul Javerzat and Hiten Madhani for generous gift of yeast strains and plasmids. We are grateful to Fabrizio Simonetti and Jocelyne Boulay for technical assistance in Northern blotting and Domenico Libri for providing reagents and analytic tools in the early phases of the project.

## Supporting information captions

**S1 Fig (related to Fig 1). The Ccr4-Not complex mediates heterochromatic gene silencing at the mating type locus. A.**, **C.**, **E.**, **G.** RT-qPCR analyses of *dg*, *tlh1*+ (**A.**), *gfp*+ (**C.**), *matMc* (**E.**) and *ura4*+ (**G.**) transcripts (mean±SD; n=3; normalized to *act1*+; relative to wt) in cells of the indicated genetic backgrounds. Two-tailed Student’s *t*-tests were used to calculate *p*-values. Individual data points are represented by black circles. **B.**, **F.** 5-fold serial diluted silencing assays using the *ade6*+ (**B.**) or *ura4*+ (**F.**) reporter genes inserted at the mating type locus (mat3M::*ade6*+; mat3M::*ura4*+). Cells of the indicated genotypes were plated on both non-selective (YES) and selective (YES low ADE or YES+5FOA) media. Defects in heterochromatic gene silencing result in the formation of white colonies (mat3M::*ade6*+) (**B.**) or sensitivity to 5FOA (mat3M::*ura4*+) (**F.**). In **F.**, *mmi1*Δ cells were also deleted for *mei4*+, since the absence of Mmi1 leads to major growth defects due to the ectopic expression of the meiosis-specific transcription factor Mei4. The mutants of interest were constructed in a *mei4*Δ background for direct comparison. **D.** Western blot showing total Gfp levels in mat3M::*gfp*+ cells of the indicated genetic backgrounds. Anti-CDC2 antibody was used as loading control.

**S2 Fig (related to Fig 2). The Ccr4-Not complex impacts heterochromatin assembly at the mating type locus. A.**, **B.** ChIP-qPCR analyses (% input; mean±SD; n=4 or 3) of histone H3 (**A.**) and TAP-tagged proteins (**B.**) in cells of the indicated genetic backgrounds. Immunoprecipitations without antibodies (no Ab) or from untagged strains were performed to determine background levels. Shown are the enrichments of *ura4*+, *dg* repeats, *tlh1*+ and *act1*+ upon immunoprecipitation with H3 antibody (**A.**) or rabbit IgG (**B.**). Individual data points are represented by black circles.

**S3 Fig (related to Fig 3). The Ccr4-Not subunits Caf1 and Mot2 regulate heterochromatin spreading at the mating type locus. A.-D.** Two-dimensional-density squarebin plots showing the red-normalized green and orange fluorescence for *caf1Δ* (**A.**, **B.**) and *mot2*Δ (**C.**, **D.**) REIII_mut_ cells grown at 32°C. A density bar represents the fraction of the most dense bin. Panels correspond to the second and third isolates for both backgrounds. **E.** RT-qPCR analyses of *cenH* transcripts (mean±SD; n=3; normalized to *act1*+; relative to wt) in cells of the indicated genetic backgrounds. Two-tailed Student’s *t*-tests were used to calculate *p*-values. Individual data points are represented by black circles.

**S4 Fig. Caf1 and Mot2 impact heterochromatin spreading at subtelomeres but not centromeres. A.**, **B.** H3K9me3 ChIP-qPCR analyses (% input; mean±SD; n=3) in cells of the indicated genetic backgrounds. Immunoprecipitations without antibodies (no Ab) were performed to determine background levels. **C.** RT-qPCR analyses of subtelomeric transcripts (mean±SD; n=3; normalized to *act1*+; relative to wt) in cells of the indicated genetic backgrounds. **A.**-**C.** Numbers correspond to the different primer pairs used in qPCR reactions and whose localization is indicated on the scheme below each graph. Two-tailed Student’s *t*- tests were used to calculate *p*-values. Individual data points are represented by black circles. In the scheme in **A.**, vertical black lines in the imr1R region represent tRNA genes that delimit heterochromatin boundaries, beyond which H3K9me3 is not enriched.

**S5 Fig (related to Fig 4). Importance of Caf1 and Mot2 catalytic activities in gene silencing and heterochromatin spreading. A.**, **D.** Western blots showing total 2xFLAG-tagged wild type or mutant Caf1 (**A.**) and Mot2 (**D.**) expressed from the pREP41 vector. Anti-CDC2 antibody was used as loading control. **B.** 5-fold serial diluted silencing assay using the *ura4*+ reporter gene inserted at the mating type locus (mat3M::*ura4*+). Cells of the indicated genotypes were plated on both non-selective (EMM-LEU) and 5FOA-containing (EMM-LEU+5FOA) media. **C.** RT-qPCR analyses of *ura4*+ transcripts (mean±SD; n=4; normalized to *act1*+; relative to wt pREP41) in cells of the indicated genetic backgrounds. Two-tailed Student’s *t*-tests were used to calculate *p*-values. Individual data points are represented by black circles.

**S6 Fig (related to Fig 5). The anti-silencing factor Epe1 opposes Ccr4-Not in gene silencing and heterochromatin spreading. A.** Northern blot showing *ura4*+ mRNA levels from total RNA samples in the indicated genetic backgrounds (mat3M::*ura4*+). The PCR probe overlapping the 3’ end of *ura4*+ also detects the endogenous *ura4-DS/E* mini-gene. BET- stained ribosomal RNAs serve as a loading control. **B.** Western blot showing total TAP-tagged Epe1 in the indicated genetic backgrounds. Anti-CDC2 antibody was used as loading control. **C.-F.** Two-dimensional-density squarebin plots showing the red-normalized green and orange fluorescence for *caf1Δ epe1Δ* (**C.**, **D.**) and *mot2*Δ *epe1Δ* (**E.**, **F.**) REIII_mut_ cells grown at 32°C. A density bar represents the fraction of the most dense bin. Panels correspond to the second and third isolates for both backgrounds.

**S7 Fig. Mutation of the Epe1 jumonji domain suppresses silencing defects in *caf1*Δ and *mot2*Δ cells. A.** Domain organization of the Epe1 protein. **B.** 5-fold serial diluted silencing assay using the *ura4*+ reporter gene inserted at the mating type locus (mat3M::*ura4*+). Cells of the indicated genotypes were plated on both non-selective (YES) and 5FOA-containing (YES+5FOA) media. **C.** RT-qPCR analyses of *ura4*+ transcripts (mean±SD; n=4; normalized to *act1*+; relative to wt) in cells of the indicated genetic backgrounds. **D.** Western blot showing total wild type or H297A TAP-tagged Epe1 in the indicated genetic backgrounds. Anti-CDC2 antibody was used as loading control. **E.** ChIP-qPCR analyses (% input; mean±SD; n=4) of the indicated strains in the mat3M::*ura4*+ background. Immunoprecipitations without rabbit IgG (no Ab) were performed to determine background levels. **C.**, **E.** Two-tailed Student’s *t*-tests were used to calculate *p*-values. Individual data points are represented by black circles.

**S1 Table. *S. pombe* strains used in this study.**

**S2 Table. Plasmids used in this study.**

**S3 Table. Oligonucleotides used in this study.**

